# Agarose floor technique: a simple scaffold-free method for 3D cell culture and multicellular tumor spheroids formation

**DOI:** 10.1101/130260

**Authors:** César Rivera

## To the editors

3D cell cultures can close the gap between *in vitro* experiments used for discovery and *in vivo* designs used for efficacy and safety assessment before proceeding to clinical studies ^1^. 3D multicellular tumor spheroids reflect better the tumor environment in terms of phenotypic heterogeneity, nutrient and oxygen gradients, intervascular domains, and micrometastases ^2^.

This letter presents the agarose floor technique to obtain single-cell suspensions from oral cancer cells in nonadherent conditions. SCC-9 cells (American Type Culture Collection, Manassas, VA, USA) were cultured in culture plastic wares with nonadhesive surface. 150 mm dish are made of nonadhesive for cells by coating with 3.2% sterile agarose (Analytical Grade, Promega, Madison, WI, USA) thin films (5-8 mL/dish). This agarose floor is allowed to dry before addition of medium (DMEM/Ham’s F12, supplemented with 10% fetal bovine serum, antibiotics and 0.4 μg/mL hydrocortisone) containing 6x10^5^ live cells/dish. The cells were maintained at 37°C in a 5% CO_2_ atmosphere and the culture medium was changed every 2 days until the sphere formation (8 days). With the passing of the days multicellular tumor spheroids are obtained. The spheres were examined with microscope and digital camera (**Figure 1**).

**Figure 1.**
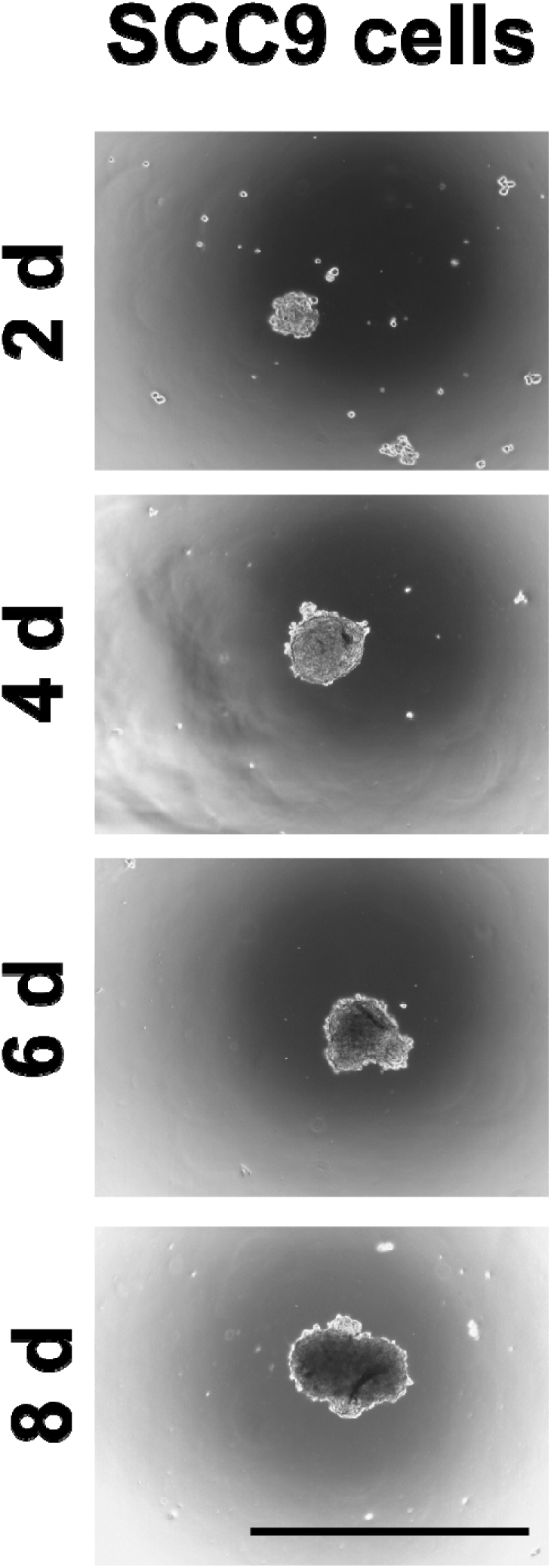
Three-dimensional multicellular tumor spheroids formation. Oral cancer cells (SCC-9) grown in non-adhesive liquid-based culture tend to form cell clusters (panels from day 2 to 8). With tumor spheroids, the cells appear fused together as “*floating popcorns in the night sky*”. Scale bar, 1mm.

Multicellular tumor spheroids are easily generated by the liquid overlay technique that prevents matrix deposition. Tumor cells are placed on tissue culture plastic covered with a thin layer of inert substrate such as agarose ^3^. Agarose is frequently used in molecular biology for the separation of large molecules, especially DNA, by electrophoresis. The pore size of a 3% agarose gel has been estimated in ∼290 nm ^4^. The low porosity, along with being an inert material, make the agarose a good candidate to obtain a non-adherent surface to epithelial cells.

## REFERENCES

[1] Zanoni M, Piccinini F, Arienti C, Zamagni A, Santi S, Polico R, Bevilacqua A, Tesei A: 3D tumor spheroid models for in vitro therapeutic screening: a systematic approach to enhance the biological relevance of data obtained. Sci Rep. 2016;6:19103.

[2] Wang J, Zhang X, Li X, Zhang Y, Hou T, Wei L, Qu L, Shi L, Liu Y, Zou L, Liang X: Anti-gastric cancer activity in three-dimensional tumor spheroids of bufadienolides. Sci Rep. 2016;6:24772.

[3] Weiswald LB, Bellet D, Dangles-Marie V: Spherical cancer models in tumor biology. Neoplasia. 2015;17(1):1–15.

[4] Pernodet N, Maaloum M, Tinland B: Pore size of agarose gels by atomic force microscopy. Electrophoresis. 1997;18(1):55–8.

